# Mimicking seasonal changes in light-dark cycle and ambient temperature modulates gut microbiome in mice under the same dietary regimen

**DOI:** 10.1101/2022.11.09.515822

**Authors:** Shoko Matsumoto, Liang Ren, Masayuki Iigo, Atsushi Murai, Takashi Yoshimura

**Author notes:** Corresponding author (TY).

## Abstract

To better adapt to seasonal environmental changes, physiological processes and behaviors are regulated seasonally. The gut microbiome interacts with the physiology, behavior, and even the diseases of host animals, including humans and livestock. Seasonal changes in gut microbiome composition have been reported in several species under natural environments. Dietary content significantly affects the composition of the microbiome, and, in the natural environment, the diet varies between different seasons. Therefore, understanding the seasonal regulatory mechanisms of the gut microbiome is important for understanding the seasonal adaptation strategies of animals. Herein, we examined the effects of changing day length and temperature, which mimic summer and winter conditions, on the gut microbiome of laboratory mice. Principal coordinate analysis and analysis of the composition of microbiomes of 16S rRNA sequencing data demonstrated that the microbiomes of the cecum and large intestine showed significant differences between summer and winter mimicking conditions. Similar to previous studies, a daily rhythm was observed in the composition of the microbiome. Furthermore, the phylogenetic investigation of communities by reconstruction of unobserved states predicted seasonal changes in several metabolic pathways. Changing day length and temperature can affect the composition of the gut microbiome without changing dietary contents.

## Introduction

The environment surrounding living organisms shows dynamic periodical changes, such as daily and seasonal fluctuations according to the Earth’s rotation. To better adapt to these periodic environmental changes, organisms have evolved endogenous biological clocks, such as the approximately 24-hour circadian clock and approximately 1-year circannual clock [1-2]. These endogenous biological clocks drive daily and seasonal rhythms in various physiological processes and behaviors.

The intestinal microbiome is known as the “microbial organ” and has been shown to be associated with physiological processes and diseases. For instance, immunoglobulin A (IgA), which is responsible for mucosal immunity, is severely reduced in germ-free (GF) animals lacking the intestinal microbiome and is restored by the administration of *Escherichia coli* [3]. Bacterial polysaccharides produced by *Bacteroides fragilis* induce CD^4+^ T-cell proliferation and are involved in the normal development of host immunity [4]. Serotonin (5-hydoxytryptamine:5-HT) is a neurotransmitter in the brain that is responsible for the regulation of various organs, including gastrointestinal peristalsis. The majority of 5-HT in the body is found in the periphery and is synthesized from intestinal chromaffin cells [5-6]. The level of 5-HT in GF- or antibiotic-treated mice is very low and has been reported to be restored by the administration of the gut microbiome [7]. The gut microbiome is not only related to immune function and the endocrine system, but it is also associated with diseases such as inflammatory bowel disease and diabetes [8].

The composition and the ratio of the gut microbiome is dynamic; it varies depending on external factors (such as diet and antibiotics) and internal factors of the host (such as age and genetic background) [9]. Recent studies have demonstrated that the composition of the gut microbiome exhibits a daily rhythm. When feces were collected every 6 hours from mice reared under conditions of 12 hours light and 12 hours dark (12L:12D), more than 60% of the detected operational taxonomic units (OTUs) of the microbiome showed daily rhythmicity. Furthermore, the deletion of circadian clock genes (e.g., *Bmal1* and *Per1/2*) results in the loss of the daily changes in the gut microbiome composition [10-11].

In addition to the daily rhythmicity, annual rhythms in compositional changes of the gut microbiome have also been reported. In a 1-year study of the Hadza people in Tanzania, the composition of the gut microbiome was different between the wet and dry seasons [12]. Analysis of fecal samples from wild gelada baboons (*Theropithecus gelada*) in Ethiopia also presented differences in gut microbiome composition between the dry and wet seasons [13]. As it is well established that food intake strongly affects the gut microbiome [14], these previous studies stated that the seasonality of the diet strongly influenced their results.

The gut microbiome is closely related to the health and diseases of host animals including poultry, livestock, and humans. Therefore, understanding the regulatory mechanisms of the gut microbiome will provide new insights into host physiology and health. To better understand the seasonal regulation of the gut microbiome, we examined the impacts of changing day length and temperature on the gut microbiome of mice, without changing the food, to evaluate the effects of seasonal cues on the gut microbiome.

## Materials and methods

### Mice and sample collection

Differences in genetic background are known to affect the gut microbiome [15]. To minimize the effects of different genetic backgrounds, inbred strains of mice are often used in microbiome studies. However, most of the commonly used inbred strains of mice, such as C57BL, DBA, BALB, 129, cannot produce melatonin due to genetic defects in the melatonin biosynthesis pathways that are important for seasonal responses [16-18]. Therefore, we used a melatonin-proficient CBA/N strain that could respond to photoperiodic changes. Ten-week-old female CBA/N mice were purchased from Japan SLC, Inc (Shizuoka, Japan). All animal experiments were approved by the Animal Experiment Committee of Nagoya University, Japan (Permit Number: A210705-001). All CBA/N mice were housed in a multi-chamber animal housing system (LP-30CCFL-8CTAR; Nippon Medical & Chemical Instruments Co., Ltd., Osaka, Japan). In this system, the light-dark cycle and temperature of each chamber can be individually controlled. Forty-eight female mice were divided into two groups and transferred to either a chamber of summer-mimicking condition (long-day and warm temperature (LW): 16 hours light and 8 hours dark, 30°C; n = 24) or a chamber of winter-mimicking condition (short-day and cool temperature (SC): 8 hours light and 16 hours dark, 10°C; n = 24). Food (Labo MR Stock, Nosan Co., Kanagawa, Japan) and water were provided *ad libitum*. After 4 weeks of exposure to each condition, four mice were euthanized using neck dislocation every 4 hours. Subsequently, the intestinal contents were collected from the cecum and large intestine (n = 4). The intestinal contents were stored at -80°C until DNA extraction.

### DNA extraction and 16S metagenomic sequencing

The total genomic DNA from each of the intestinal contents was extracted following the standard protocol according to the instructions of ISOSPIN Fecal DNA (315-08621; NIPPON GENE Co., Ltd., Tokyo, Japan). DNA concentration was quantified using a Qubit® 3.0 Fluorometer (Q33216; Thermo Fisher Scientific, Waltham, MA, USA) and a Qubit™ dsDNA HS Assay Kit (Q32851; Thermo Fisher Scientific). The 16S metagenomic sequencing was performed according to the procedure set by Illumina (San Diego, CA, USA; https://support.illumina.com/documents/documentation/chemistry_documentation/16s/1 6s-metagenomic-library-prep-guide-15044223-b.pdf) with a few modifications. The first PCR amplified the V3–V4 region of the bacterial 16S rRNA gene, which used 16S Amplicon PCR Forward Primer (5′-TCGTCGGCAGCGTCAGATGTGTATAAGAGACAGCCTACGGGNGGCWGC AG-3′) and 16S Amplicon PCR Reverse Primer (5′-GTCTCGTGGGCTCGGAGATGTGTATAAGAGACAGGACTACHVGGGTATC TAATCC-3′). The first PCR reactions were carried out in a total volume of 10 μL that consisted of 5.0 μL KAPA HiFi HotStart ReadyMix (KK2602; KAPA Biosystems, Potters Bar, Herts., UK.), 0.25 μL each primer (10 pmol/L), 0.25 μL SYTO9 (25 μmol/L; S34854; Thermo Fisher Scientific), 5 ng of DNA template, and pure water. Pre-denaturation was conducted at 95°C for 5 min, followed by denaturation at 95°C, annealing at 55°C, and extension at 72°C for 30 s. These three steps were repeated for 26 cycles and were finally extended to 72°C for 5 min. The PCR products were cleaned with a 1:1 ratio of SeraPure Magnet Beads [19] to PCR-amplified DNA, and the amplicon concentration was measured using the Qubit system. A second round of PCR amplification was performed to add an index primer for the barcode sequence. The second PCR reactions were carried out in a total volume of 10.0 μL that consisted of 5.0 μL KAPA HiFi HotStart ReadyMix (KAPA Biosystems), 25 μmol/L of SYTO9 (Thermo Fisher Scientific), 1.75 μL pure water, 1.0 μL of the first PCR product, and 1.0 μL of each index primer of the Nextera XT Index Kit v2 Set B and Set C (FC-131-2002, FC-131-2003; Illumina Inc., CA., USA.). The first step, redenaturation, was conducted at 95°C for 5 min and allowed to run for eight cycles; the other cycle steps were the same as in those in the first PCR. The second PCR products were cleaned with a 7:10 ratio of SeraPure Magnet Beads to PCR-amplified DNA, and the amplicon concentration was measured again using the Qubit system. The PCR products of each sample were mixed to prepare PCR amplicon libraries. Quality checks of each library were conducted using a Bioanalyzer 2100 (Agilent Technologies Japan, Ltd., CA., USA.) and an Agilent DNA 1000 Kit (5067-1504; Agilent Technologies Japan, Ltd.). After quality assessment and quantification, the same amounts of amplified products were collected and sequenced using paired-end 2 × 300 bp in MiSeq (Illumina).

### Bioinformatic Analyses

The 16S rRNA amplicon sequence data have been deposited in the Sequence Read Archive (SRA) under the project ID PRJNA874855 (www.ncbi.nlm.nih.gov/sra). The resulting data were analyzed using Quantitative Insights Into Microbial Ecology 2 (QIIME2) (https://docs.qiime2.org/2019.7/install/ version 2019.7 or 2020.6), GraphPad Prism, and R. After trimming low-quality bases from the demultiplexed reads, overlapping paired-end reads were merged and the sequencing data were denoised using the Divisive Amplicon Denoising Algorithm 2 (DADA2) in QIIME2. The low-quality parts of the sequences from the forward and reverse reads were removed by trimming to 220 and 180 bases. The forward and reverse reads were then merged and the chimeric sequences were removed. Samples with more than 20,000 reads were used for the analyses. The Silva classifier (https://www.arb-silva.de/documentation/release-138/ ver.138) was used as the reference template at a 99% similarity level.

### Statistical analysis

Principal coordinate analysis (PCoA) is a general method of dimensionality reduction that can choose measures other than Euclidean distance, unlike principal component analysis (PCA), which uses Euclidean distance. We performed PCoA using the qiime2R library in R. Permutational multivariate analysis of variance (PERMANOVA) was performed to detect groups that differed significantly between intestinal sites, seasonal conditions, and sampling time points. PERMANOVA is used to examine whether there is a significant difference between bacterial flora when focusing on a certain parameter [20]. PERMANOVA was performed according to the QIIME2 protocol using four scales: the Bray–Curtis distance, unweighted/weighted UniFrac distance, and Jaccard distance matrix; and the number of permutations was set to 999. From the Bray–Curtis distance, significant differences were detected in all parameters, such as intestinal sites, seasonal conditions, and sampling time. JTK_Cycle is a program that detects rhythmicity using time-series data [21]. The JTK_Cycle was performed using the R’s MetaCycle library. The Student’s *t*-test or Kruskal–Wallis test was performed using GraphPad Prism to compare two samples. The analysis of composition of microbiomes (ANCOM) is a statistical method for comparing the microbiome composition of two or more groups [22]. ANCOM was performed using QIIME2 to clarify whether the microbiome differed significantly under different seasonal conditions. The phylogenetic investigation of communities by reconstruction of unobserved states (PICRUSt) is a method that predicts functional composition by collating sequence results and a database of functions [23]. PICRUSt was performed using QIIME2 to predict the gene function of bacterial flora.

## Results

Ten-week-old female CBA/N mice were transferred to long-day and warm temperature (LW) summer-mimicking conditions (16 hours light / 8 hours dark, 30°C) or short-day and cool temperature (SC) winter-mimicking conditions (8 hours light / 16 hours dark, 10°C). After 4 weeks of exposure to LW or SC, the intestinal contents were collected from the cecum and large intestine every 4 hours (n = 4). The gut microbiome was determined via 16S ribosomal RNA (rRNA) sequencing using the time-series samples collected under SC and LW conditions. Beta diversity quantifies (dis)similarities between samples. The most popular beta diversity measures in microbiome research include the Bray–Curtis index, Jaccard index, and UniFrac distances. When we performed PCoA using the the Bray–Curtis distance, unweighted/weighted UniFrac distance and Jaccard distance matrices, all measures presented significant differences between the cecum and large intestine (PERMANOVA, Bray–Curtis distance *q* = 0.001; Unweighted UniFrac distance *q* = 0.001; weighted UniFrac distance *q* = 0.001; Jaccard distance matrix *q* = 0.041). This suggests that the microbiome from the cecum and large intestine forms significantly different clusters (Fig 1A).

**Fig 1.**
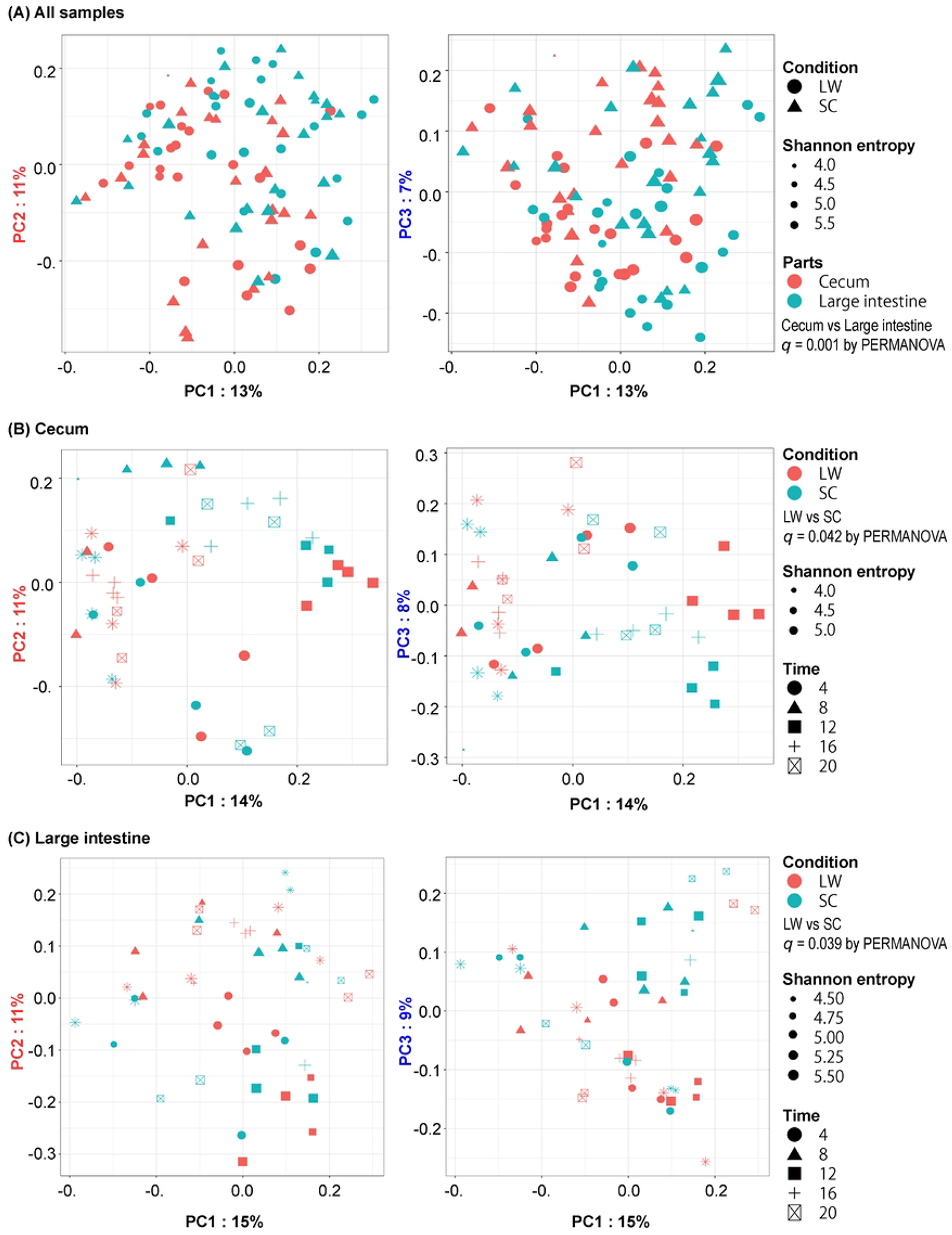
Differences in the microbiome under summer- and winter-mimicking conditions. Mouse gut microbiome forms different clusters depending on seasonal conditions and sampling time. Principal coordinate analysis (PCoA) plot with Bray Curtis distance in the gut microbiome of CBA/N mice. (A) PCoA plot using all samples of long day and warm (LW) and short day and cool (SC) groups. Each point represents one sample. The shape of the dot represents the seasonal conditions. The size of the dot represents Shannon diversity, and the color of the dot represents the intestinal sites (Cecum, Large intestine). (b and c) PCoA plots in the cecum (B), and the large intestine (C). The color of the dots indicates the seasonal conditions (LW, SC). The size of the dots indicates the Shannon diversity, and the shape of the dots indicates the sampling time (0:00, 4:00, 8:00, 12:00, 16:00, 20:00). The number of individuals at each sampling time was 4. However, the number was 2 at two points (cecum LW 8:00 and large intestine SC 16:00) because some samples failed in the quality control process.

Because the microbiome from the cecum and large intestine formed different clusters, we then examined the PCoA for the cecum and large intestine separately. A PERMANOVA test demonstrated significant differences between summer- and winter-mimicking conditions in the cecum (Fig 1B) [Bray–Curtis distance (*p* = 0.042) and Jaccard distance (*p* = 0.009)] and in the large intestine (Fig 1C) [Bray–Curtis distance (*p* = 0.039) and Weighted UniFrac distance (*p* = 0.035)]. To show the differences between the two seasonal conditions, the results of Figs 1B and 1C were re-plotted using only the seasonal conditions (S1 Fig). The PERMANOVA test also demonstrated significant differences among different times of the day for both the cecum (Fig 1B, S1 Table) and the large intestine (Fig 1C, S2 Table). These results suggest that the flora of both the cecum and large intestine of CBA/N mice form different clusters depending on the seasonal conditions and times of the day.

As significant differences were observed at different times of the day (Fig 1; S1 and S2 Tables), daily variations in the relative abundance of the microbiome at the phylum level was analyzed under summer- and winter-mimicking conditions (Fig 2). Bacteroidota and Firmicutes were dominant in all samples, followed by Deferribacterota and Desulfobacterota (Figs 2A and 2C). Because the results of Figs 2A and 2C suggest possible daily variations in the composition of bacterial flora in the cecum and large intestine, daily rhythmicity in the relative abundance at the phylum level was examined using the JTK_Cycle incorporated in the MetaCycle package in R [21]. This analysis demonstrated that Bacteroidota and Deferribacterota under LW condition, and Desulfobacterota under SC conditions, showed clear daily rhythmicity in the cecum, whereas Desulfobacterota showed daily rhythmicity under both LW and SC conditions in the large intestine (*p* < 0.05). Because the bar plots showed daily changes at both intestinal sites, data from the middle of the day (12:00) and middle of the night (0:00) were also compared (S2 Fig), confirming clear differences in the composition of the microbiome between winter- and summer-mimicking conditions.

**Fig 2.**
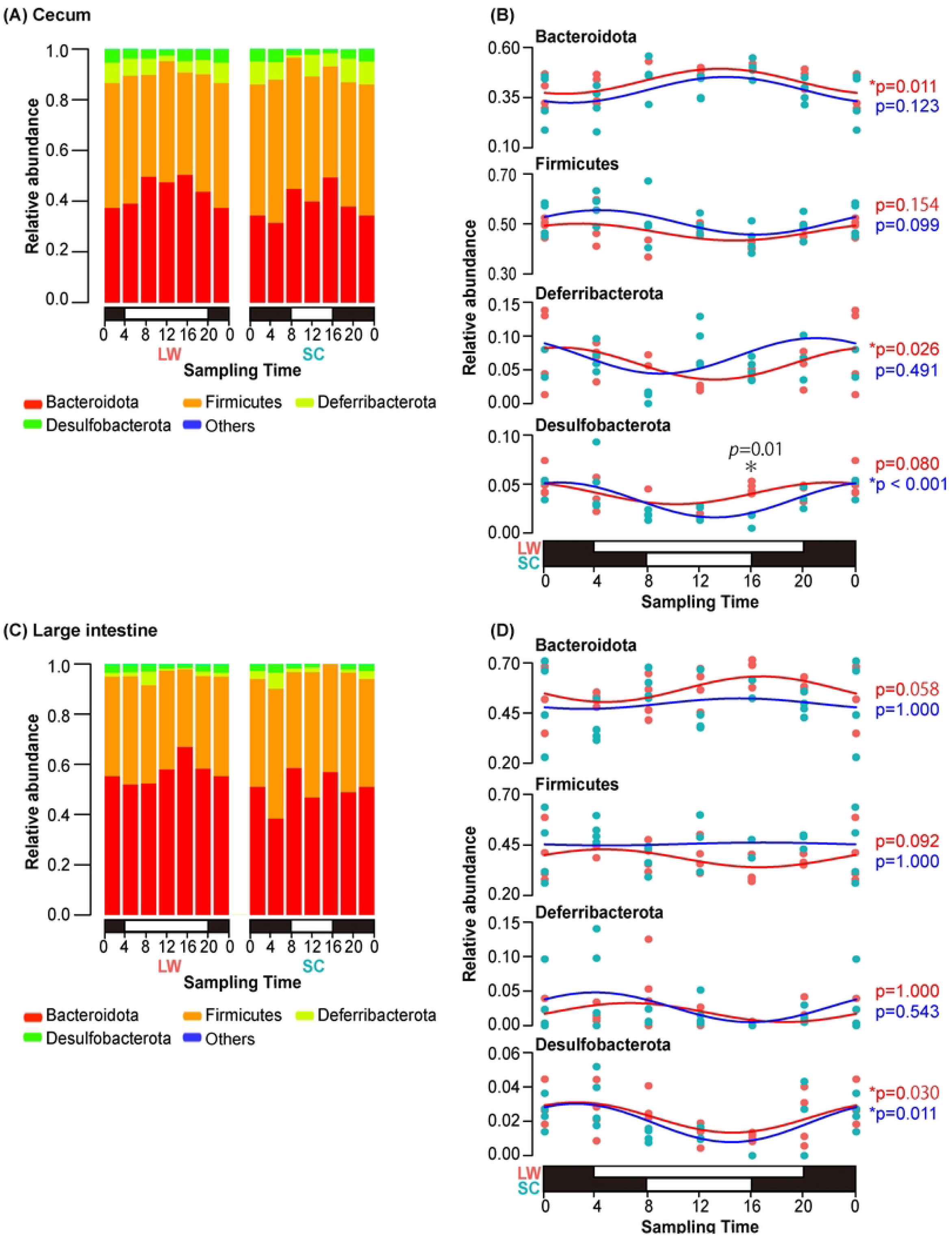
Daily variation in the composition of the microbiome under summer- and winter-mimicking conditions. Relative abundance of phylum level in (A) cecum and (C) large intestine of CBA/N mice. The horizontal axis indicates the sampling time, and the vertical axis indicates the average relative abundance of each bacterial phylum. The white and black bar with the horizontal axis indicates the light/dark cycle under LW/SC conditions. The number of individuals at each sampling time was 4. However, the number was 2 at two points (cecum LW 8:00 and large intestine SC 16:00) because some samples failed in the quality control process. The data of 0:00 were double-plotted for the same data. Curve fitting of the major bacterial phyla in (B) the cecum and (D) the large intestine and their relative abundance. Each dot represents one sample, and the color indicates the seasonal condition (red: LW; blue: SC). *P*-values of the JTK_Cycle are on the right side of the graph. Asterisk indicates significant difference detected by JTK_Cycle or *t*-test between LW and SC conditions.

We then compared the composition of microbiomes under LW and SC conditions, at the phylum and genus levels, by using data from all sampling points to analyze the composition of microbiomes (ANCOM) (Fig 3). The comparison at the phylum level revealed that the relative abundance of Bacteroidota was significantly higher in the LW than in the SC (Fig 3A, S3 Table). Meanwhile, a genus level comparison detected significant differences in four genera (*Lactobacillus, Bacteroides, Alloprevotella*, Muribaculaceae; S3 Fig, S3 Table) in the cecum. Although no significant difference was observed in the large intestine (S4 Table), the ratio of Firmicutes to Bacteroidota (F/B ratio) was significantly higher in the winter-mimicking conditions than in the summer-mimicking conditions in both the cecum and the large intestine (Fig 3B) (*t*-test, cecum *p* = 0.044; large intestine *p* = 0.011).

**Fig 3.**
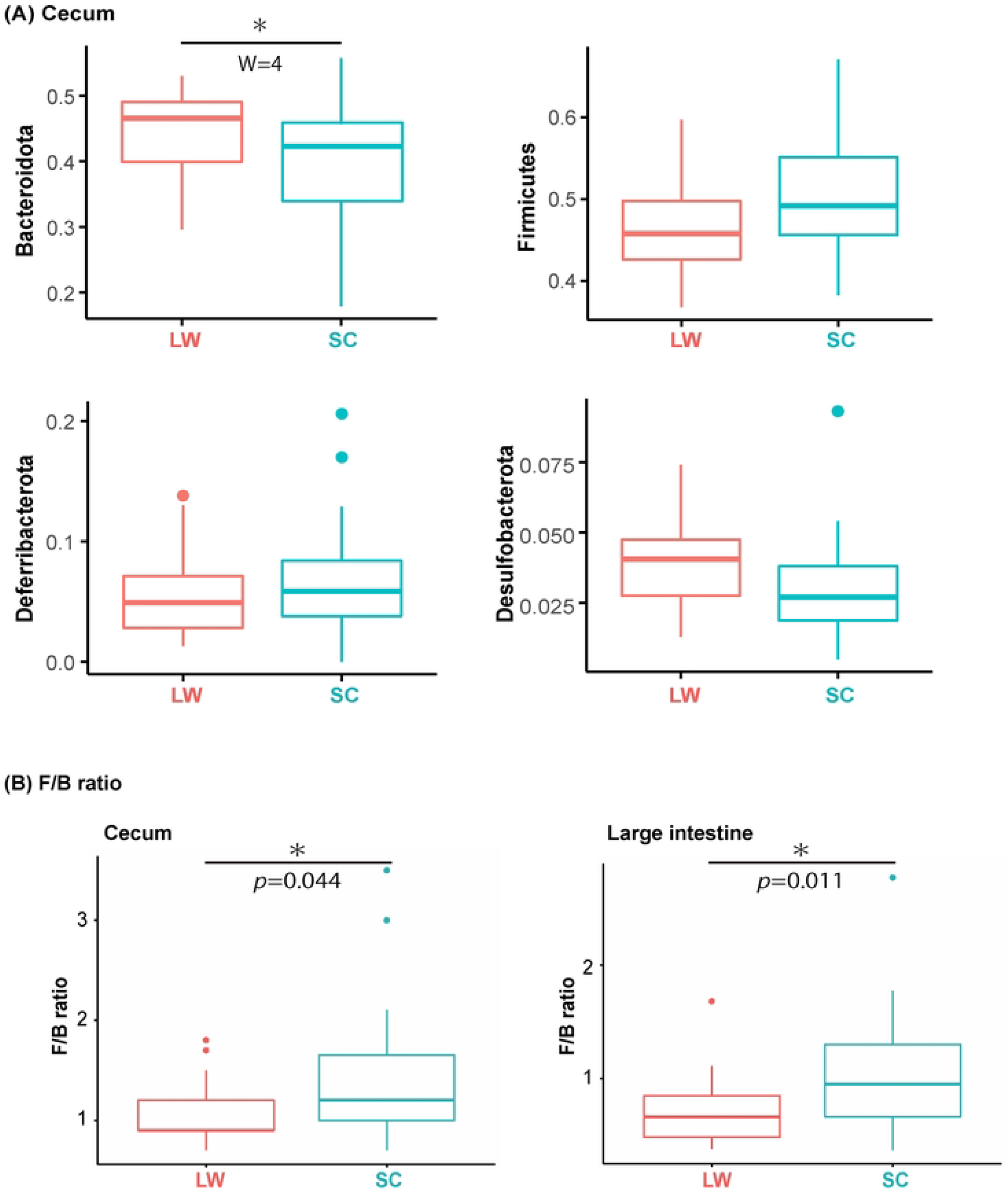
Bacteroidota in the cecum differ significantly between summer- and winter-mimicking conditions. (A) Relative abundance of Bacteroidota in the cecum is different between summer- and winter-mimicking conditions. Analysis of composition of microbiomes (ANCOM) was performed on the four major phyla that constitute the cecum and large intestine flora of CBA/N mice. Boxplots showing bacterial genera with significantly different relative abundances depending on seasonal conditions (LW, SC) (cecum, LW n = 22; SC n = 24). Plots show medians and interquartile ranges. W represents the number of null hypotheses rejected when statistics are performed to see if there is a difference between samples for a particular bacterium. (B) The Firmicutes/Bacteroidota (F/B) ratios in cecum (left) and large intestine (right) were compared between LW and SC conditions, respectively. Box plots show medians and interquartile ranges, and symbols indicate the significant differences by *t*-test (*; *p* < 0.05).

Beta diversity quantifies (dis-)similarities between samples, whereas alpha diversity is an indicator used to assess the distribution of species abundance in a given sample. Therefore, we compared alpha diversity values using the Kruskal–Wallis test (Fig 4). Using the observed feature scale, which indicates the number of taxa observed, a significant difference between LW and SC was observed in the cecum (Fig 4B). Using Pielou’s evenness index, which weighs the evenness of the groups, a significant difference between LW and SC was observed in the large intestine (Fig 4C).

**Fig 4.**
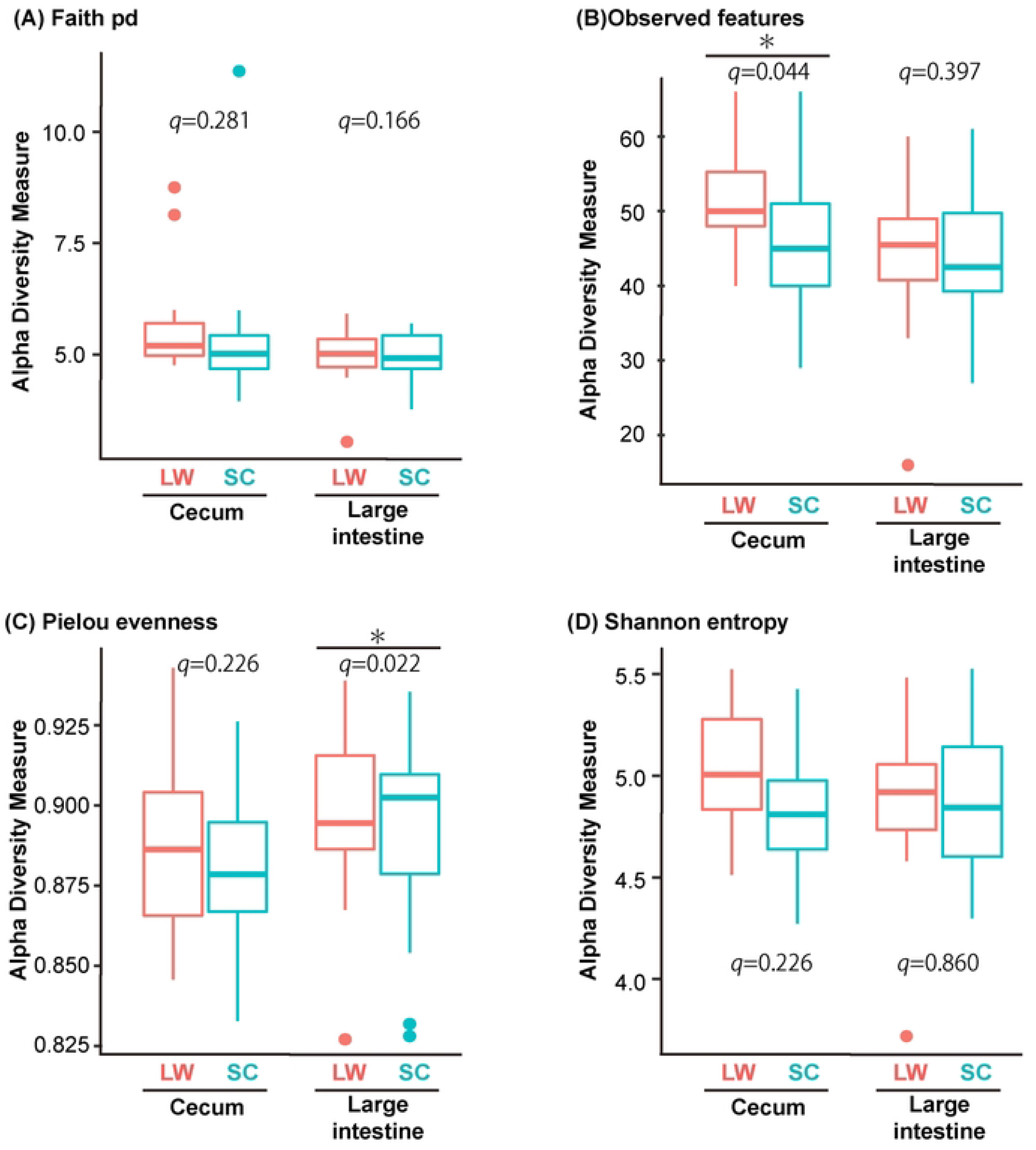
Species abundances are slightly higher under LW conditions than under SC conditions. Difference in alpha diversities in the cecum and the large intestine between summer- and winter-mimicking conditions. Alpha diversity indices used were Faith pd, Observed features, Pielou evenness and Shannon entropy. Diversities that are significantly different by the Kruskal-Wallis test are indicated by asterisks (* *q* < 0.05).

PICRUSt is a computational approach to predict the functional composition of a metagenome using marker genes, such as the 16S rRNA gene, and a database of reference genomes [23]. PICRUSt has been shown to recapture key findings from the Human Microbiome Project, and accurately predict the abundance of gene families in host-associated and environmental communities. Therefore, we used PICRUSt to estimate the changes in metabolic pathways in our analysis (Fig 5). Genes for which significant differences were determined by statistical analysis using ANCOM are shown in Fig 5, with focus on changes in the metabolic pathways associated with daily variations in the intestinal microbiome in the cecum (S3 Table) and large intestine (S4 Table). Many of the metabolic pathways, for which significant differences were detected, presented increases during the day (i.e., 12:00 and 16:00) when sleeping (Fig 5). When these pathways were mapped using KEGG mapper, multiple metabolic pathways were detected under LW conditions in the cecum (Fig 5A). In contrast, significant differences in CRISPR system cascade subunits were detected in both the cecum (Fig 5B) and the large intestine (Fig 5D) under SC conditions.

**Fig 5.**
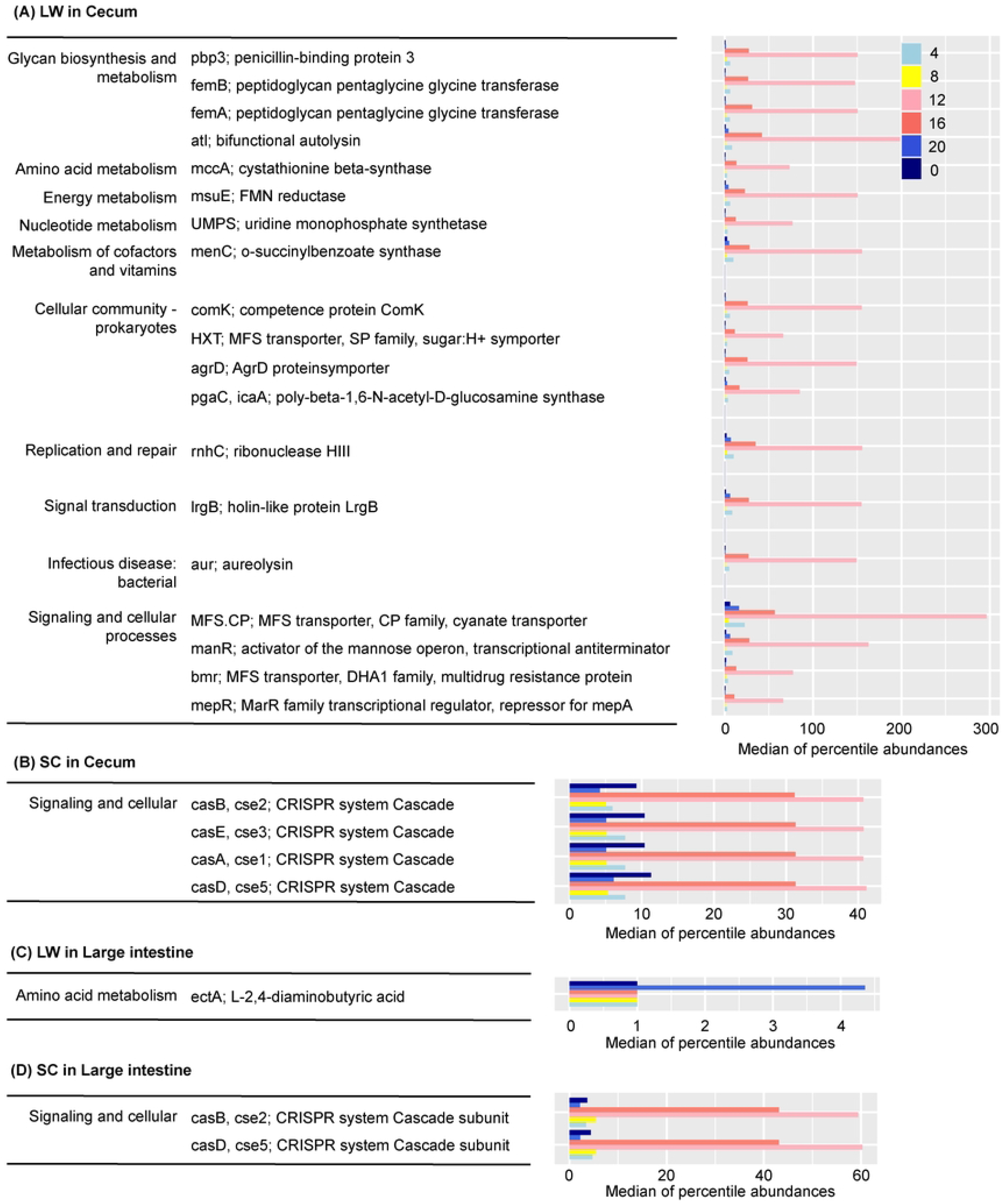
Pathways predicted to show daily changes. Predicted daily changes in the KEGG pathways in the cecum under (A) LW and (B) SC conditions. Predicted daily change in the KEGG pathways in the large intestine under (C) LW and (B) SC conditions. PICRUSt prediction results that focused on daily changes of gut microbiome in cecum and large intestine of CBA/N mice. The left legend represents the KEGG pathways determined by ANCOM to be significantly different by sampling time (left; level 2, right; level 6). The bar graph on the right represents the median of percentile abundances each functional pathway.

Thereafter, we selected genes for which ANCOM predicted to present significant differences between the LW and SC conditions (Fig 6). This analysis detected 29 genes in the cecum, but no genes were detected in the large intestine. When these genes were mapped to KEGG pathways using KEGG mapper, enrichment in the metabolism of vitamins, cofactors, amino acids, and carbohydrates was observed under LW conditions, compared with that under SC conditions (Fig 6).

**Fig 6.**
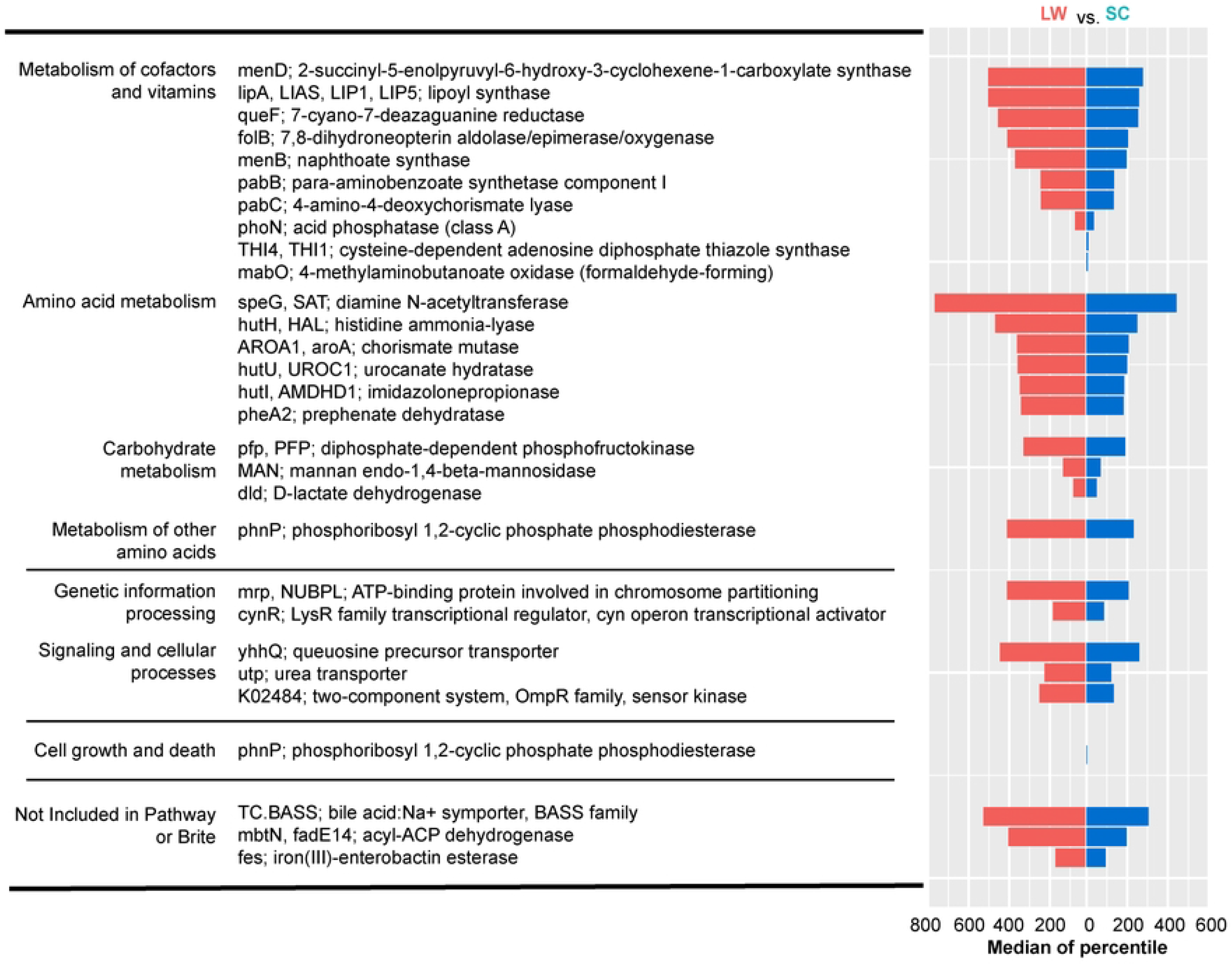
Pathways predicted to differ between LW and SC conditions. PICRUSt prediction results that focused on seasonal conditions of cecum microbiome in CBA/N mice. The left legend represents the KEGG pathways determined by ANCOM to be significantly different by seasonal condition (left; level 2, right; level 6). The bar graph on the right represents the median of percentile abundances each functional pathway.

## Discussion

The gut microbiome is closely related to the physiology of, and the diseases affecting, host animals [3, 6, 8]. Seasonal changes in the microbiome have been reported in humans and baboons living under natural environmental conditions. These seasonal changes are considered to the difference in the diet between dry and wet seasons [12-13]. The aim of this study was to investigate whether changes in day length and temperature can cause changes in the intestinal microbiome without changing dietary contents under experimental conditions.

Consistent with previous studies [24], the microbiomes of the cecum and large intestine formed significantly different clusters (Fig 1). The major phyla constituting the bacterial flora in the cecum and large intestine of the CBA/N mice were Bacteroidota, Firmicutes, Deferribacterota, and Desulfobacterota (Fig 2). Although differences in genetic background are known to affect microbiome composition, these four phyla are also dominant in the intestinal tract of C57BL/6J mice, a common laboratory mouse strain [25]. Bacteroidota and Firmicutes are common members of the intestinal flora in many organisms, including humans. Bacteroidota are primarily involved in the breakdown of proteins and carbohydrates. They are also responsible for T-cell-mediated immune responses and maintenance of the intestinal environment through butyrate production [26]. Similarly, Firmicutes are also involved in the breakdown of carbohydrates, and are a fundamental part of life support in mice [27]. In contrast, Deferribacterota is a mouse-specific phylum that is not found in human intestinal flora. Recent studies have reported daily variations in the intestinal microbiome of mice [10, 28]. In line with these reports, we confirmed the daily variations in the composition of bacterial phyla in the present study (Fig 2). In addition to the daily variations, the PERMANOVA test demonstrated significant differences between the summer- and winter-mimicking conditions in the cecum and large intestine. Furthermore, ANCOM demonstrated that the relative abundance of the phylum Bacteroidota in the cecum was significantly higher under summer-mimicking conditions (Fig 3). The F/B ratio of CBA/N mice was significantly increased in winter-mimicking conditions in both the cecum and large intestine (Fig 3). An increase in the F/B ratio in winter has been reported in musk deer (*Moschus* spp.) and Tibetan macaques (*Macaca thibetana*) [29-30], its increase is reported summer in human [31]. Thus, the patterns of seasonal change vary among species. However, it is interesting to note that cold exposure has been reported to increase the proportion of Firmicutes and, hence, the F/B ratio in mice [32].

Alpha diversity summarizes the distribution of species abundances, and microbiome from summer-mimicking conditions showed higher diversity than those from winter-mimicking conditions for most of the examined parameters (Fig 4). PICRUSt predicted the abundance of gene families in host-associated and environmental communities. Interestingly, multiple metabolic pathways were detected under LW conditions in the cecum using PICRUSt (Fig 6). Nevertheless, further studies are needed to understand the functional significance of these seasonally regulated metabolic pathways.

In conclusion, this study clearly demonstrated that seasonal factors of the environment, such as day length and temperature, can change the composition of the gut microbiome without changes in the dietary content. Nevertheless, how these seasonal changes affect the physiology, behavior, and diseases of host animals requires elucidation in future studies.

## ACKNOWLEDGEMENTS

This work was supported by a JSPS KAKENHI Grant-in-Aid for Scientific Research (S) (19H05643) and JST SPRING, grant number JPMJSP2125(RB201019). The author would also like to take this opportunity to thank the Interdisciplinary Frontier Next-Generation Researcher Program of the Tokai Higher Education and Research System.

## Supporting information

**S1 Fig. Mouse gut microbiome forms different clusters depending on seasonal conditions**. Principal coordinate analysis (PCoA) plot with Bray Curtis distance in (A) the cecum and (B) the large intestine microbiome of CBA/N mice. Each point represents one sample, and the color of the dots indicates the seasonal conditions (LW, SC). The number of individuals at each sampling time was 4. However, the number was 2 at two points (cecum LW 8:00 and large intestine SC 16:00) because some samples failed in the quality control process.

**S2 Fig. Relative abundance at phylum level**. Relative abundance at phylum level in (A) the cecum and (B) the large intestine of CBA/N mice at day and night. Day is the data from the middle of the day (12:00), while night is the data from the middle of the night (0:00). The number of individuals at each bar is 4.

**S3 Fig. Relative abundance at genus level is different between summer- and winter-mimicking conditions in the cecum**. Analysis of composition of microbiomes (ANCOM) detected the significant differences between LW and SC in Lactobacillus, Bacteroides, Alloprevotella and Muribaculaceae. W represents the number of null hypotheses rejected when statistics are performed to see if there is a difference between samples for a particular bacterium. The number of LW individuals is 22, and SC individuals is 24.

**S1 Table. Results of PERMANOVA in the cecum of CBA/N mice**. The beta diversity measure used is (A) Bray Curtis distance (B) Unweighted unifrac distance (C) Weighted unifrac distance and (D) Jaccard distance matrix.

**S2 Table. Results of PERMANOVA in the large intestine of CBA/N mice**. The beta diversity measure used is (A) Bray Curtis distance (B) Unweighted unifrac distance (C) Weighted unifrac distance and (D) Jaccard distance matrix.

**S3 Table. Results of ANCOM in phylum and genus level in the cecum of CBA/N mice**. The left side shows the results at the gate level and the right side shows the results at the genus level. W indicates the number of rejected hypotheses, and the Reject null hypothesis column indicates TRUE for those judged significant by the automatically set threshold.

**S4 Table. Results of ANCOM in phylum and genus level in the large intestine of CBA/N mice**. The left side shows the results at the gate level and the right side shows the results at the genus level. W indicates the number of rejected hypotheses, and the Reject null hypothesis column indicates TRUE for those judged significant by the automatically set threshold.

